# MuDoGeR: Multi-Domain Genome Recovery from metagenomes made easy

**DOI:** 10.1101/2022.06.21.496983

**Authors:** Ulisses Nunes da Rocha, Jonas Coelho Kasmanas, René Kallies, Joao Pedro Saraiva, Rodolfo Brizola Toscan, Polonca Štefanič, Marcos Fleming Bicalho, Felipe Borim Correa, Merve Nida Baştürk, Efthymios Fousekis, Luiz Miguel Viana Barbosa, Julia Plewka, Alexander J. Probst, Petr Baldrian, Peter F. Stadler, CLUE-TERRA consortium

## Abstract

Several computational frameworks and workflows that recover genomes from prokaryotes, eukaryotes, and viruses from metagenomes exist. However, it is difficult for scientists with little bioinformatics experience to evaluate quality, annotate genes, dereplicate, assign taxonomy and calculate relative abundance and coverage of genomes belonging to different domains. MuDoGeR is a user-friendly tool accessible for non-bioinformaticians that make it easy to recover genomes of prokaryotes, eukaryotes, and viruses from metagenomes, either alone or in combination. We tested MuDoGer using 24 individual-isolated genomes and 574 metagenomes, demonstrating the applicability for a few samples and high throughput. MuDoGeR is open-source software available at https://github.com/mdsufz/MuDoGeR.

## Introduction

Metagenomics encompasses the concepts and techniques used to study genetic material recovered directly from mixed microbial communities (Dias et al., 2019; Keller-Costa et al., 2021; López-Mondéjar et al., 2020; Oliveira Monteiro et al., 2022; J. P. Saraiva et al., 2017; J. P. Saraiva, Worrich, et al., 2021; Tláskal et al., 2021; Wooley & Ye, 2010). High-throughput sequencing reads have been used to reassemble genomes since 2004 (Tyson et al., 2004), although the exponential increase in genome recovery only occurred after 2011 with the development of many specialized metagenomic assemblers (Koren et al., 2011; Li et al., 2016; Namiki et al., 2012; Nurk et al., 2017; Peng et al., 2012). In 2013, differential coverage binning was used in the first open-source tool for genome-resolved metagenomics (Albertsen et al., 2013; Sharon et al., 2013). Since then, the bioinformatics community has developed different frameworks specific to genome recovery of prokaryotes (Corrêa et al., 2020; Sieber et al., 2018; Uritskiy et al., 2018), eukaryotes (Corrêa et al., 2020; Kasmanas et al., 2021; West et al., 2018), and viruses (Corrêa et al., 2020; Guo et al., 2021; Kallies et al., 2019; Kieft et al., 2020; Ren et al., 2017). Genome-centric metagenomics allows the simultaneous exploration of individual populations’ functional potential and phylogeny of uncultivated species(Evans Paul N. et al., 2015; Liu et al., 2022; Melkonian et al., 2021; J. P. Saraiva, Bartholomäus, et al., 2021; J. P. L. F. Saraiva et al., 2017; Tláskal et al., 2021). Although software developers created several genome-centric tools, most require users with expertise in bioinformatics or computational biology to understand the overall pipeline, install the different dependencies, run a complex framework that is often written in various computer languages, and understand the output. To extend the use of genome-centric metagenomics to those with entry-level expertise in bioinformatics and assure the data generated can be reused, the next steps in software development in the field need to guarantee that (i) the tools can be used, installed, and maintained by non-experts in computational biology and bioinformatics (Mangul et al., 2019) and (ii) data generated by any tool follow the FAIR guiding principles (Corrêa et al., 2020; Kasmanas et al., 2021; Wilkinson et al., 2016).

Here we present MuDoGeR, a framework that makes it easy to recover genomes from prokaryotes, eukaryotes, and viruses from metagenomes. We developed MuDoGeR to be easily installed and usable by non-experts. Our modular framework allows one to use the same input to recover genomes from prokaryotes, eukaryotes, and viruses alone or in combination. At the same time, users may install and use the individual modules independently. We also constructed our framework to ensure the outputs could be used to simultaneously study the genetic potential and phylogeny of the individual species from which the genomes were recovered.

## Methods

MuDoGeR was divided into five modules (Figure 1A). Module 1 deals with the pre-processing of the raw sequences. After module 1, MuDoGeR is divided into three branches (Modules 2-4). Module 2 can be used to recover prokaryotic metagenome-assembled genomes (MAGs). Module 3 was assembled to generate uncultivated viral genomes (UViGs), and Module 4 can retrieve eukaryotic metagenome-assembled bins (MABs). We recover eukaryotic MABs as eukaryotes usually need much higher coverage to assemble than prokaryotic MAGs, allowing those interested in eukaryotes to study even partial eukaryotic genomes. Module 5 was designed to generate outputs to be used in genome-centric biodiversity analysis, and users can use it to calculate the coverage and relative abundance table of the recovered MAGs, UViGs, MABs, and Open Reading Frames (ORFs) in Modules 2-4.

**Figure 1.**
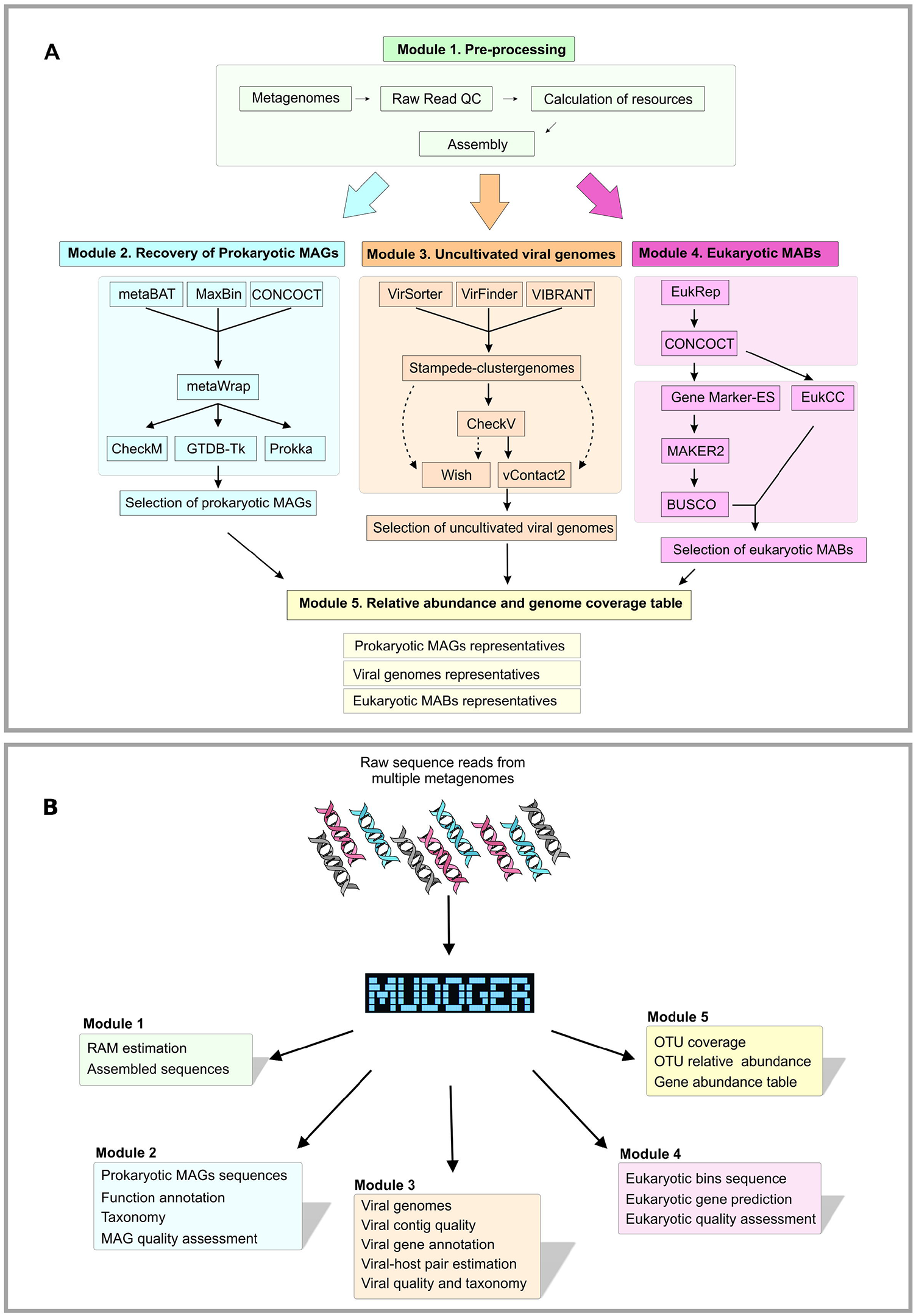
MuDoGeR workflow and outputs. **A -** MuDoGeR modular workflow. Module 1 – Pre-processing; Module 2 – Recovery of prokaryotic metagenome-assembled genomes (MAGs); Module 3 – Recovery of Uncultivated viral contigs (UViGs); Module 4 – Recovery of eukaryotic metagenome-assembled bins (MABs); Module 5 – Calculation of coverage and relative abundances. **B -** MuDoGeR modular outputs. In module 1, Random Access Memory (RAM) is calculated (for optimization purposes), and sequences are assembled. In module 2, all prokaryotic MAG sequences are stored, assigned taxonomy, annotated, and quality assessed. In module 3, all viral contigs are stored, their quality assessed, taxonomy assigned, and annotated. Additional outputs include the estimation of viral hosts. Outputs of module 4 include the storage of all eukaryotic bin sequences, quality assessment, and gene prediction. Module 5 outputs consist of tables containing the Operation Taxonomic Units (OUT) coverage, relative abundances, and gene abundance.

Currently, MuDoGeR works on paired-end short-sequence reads generated by ILLUMINA machines, but future updates will include tools to work with data from long-read sequencing. The MuDoGeR framework is a wrapper of more than 20 tools and 50 custom scripts written using Shell, Python, and R. It was designed to be an easy-to-use tool that outputs ready-to-use comprehensive files (Figure 1B). After installation, the user can run the complete pipeline using five commands, one for each module. Each module of MuDoGeR generates several outputs that users may use genome-centric analysis focusing on genetic potential, phylogenetic biodiversity of MAGs, UViGs, and MABs individually or merge both types of analysis in the same study (Figure 1B).

### MuDoGeR can be easily installed

Part of the effort to make MAGs recovery “easy” involves making the complex bioinformatics tools easily accessible, requiring little user input. A MAG/UViG/MAB recovery pipeline requires several tools that have multiple dependencies. More often than not, these tools have conflicting dependencies. Consequently, making them cooperate in a single operating system is challenging even for bioinformatics and computational biology experts. To tackle this problem, MuDoGeR uses the Conda environment management system (Grüning et al., 2018) to create multiple environments automatically orchestrated during the pipeline’s functioning. The installation protocol creates and automatically sets up 24 independent environments.

In addition, several MAG/UViG/MAB recovery tools require specific databases to work. Knowing the correct database and adequately installing them can also limit the genome recovery process by non-bioinformaticians. In MuDoGeR, we also developed a database “download and set up”-tool that makes them ready to use. Currently, MuDoGeR makes 11 databases available and integrated into our pipeline.

### MuDoGeR v1.0 at a glance

The only mandatory inputs for MuDoGeR are the paths to the samples’ raw sequencing reads. Following, the user can use one command to run each complete module. Module 1 starts by using the procedure implemented in metaWRAP (Uritskiy et al., 2018) to quality control the raw sequences. Following this, we created a regression model to estimate the Random Access Memory (RAM) required to assemble metagenomes with metaSPAdes (Nurk et al., 2017) using the 574 metagenomic libraries. From these, 558 (97.2%) fell in the predicted range (Supplementary Figure 1). The total RAM correlates highly with chimer frequencies (Supplementary Figure 2). MuDoGeR then uses metaSPAdes for assembly.

Module 2 integrates the prokaryotic binning procedure implemented in metaWrap using Metabat2 (Kang et al., 2015), Maxbin2 (Wu et al., 2016), and CONCOCT (Alneberg et al., 2014) binning tools. MuDoGeR uses CheckM (Parks et al., 2015) and GTDB-tk (Chaumeil et al., 2020) to estimate quality and taxonomy. Moreover, Prokka (Seemann, 2014) annotates ORFs, and BBtools (sourceforge.net/projects/bbmap/) is employed to calculate sequence metrics from the prokaryotic MABs. Finally, MuDoGeR summarizes all the outputs and selects prokaryotic MAGs as defined by Parks et al. (2018) (Parks et al., 2017).

The recovery of UViGs, performed in module 3, starts by integrating the viral sequence recovery tools VirSorter2 (Guo et al., 2021), VirFinder (Ren et al., 2017), and VIBRANT (Kieft et al., 2020). Later, the potential viral sequences are dereplicated using Stampede-clustergenomes (https://bitbucket.org/MAVERICLab/stampede-clustergenomes/src/master/). Subsequently, putative viral sequences are annotated using vConTACT2 (Bin Jang et al., 2019), and quality assessment is performed using CheckV (Nayfach et al., 2021). MuDoGeR also uses WIsH (Galiez et al., 2017) to predict the potential prokaryotic hosts from the recovered viral sequences by integrating the results from modules 2 and 3. Finally, MuDoGeR compiles the tools’ outputs and selects high-quality and complete UViGs as indicated by CheckV developers (Nayfach et al., 2021).

In Module 4, MuDoGeR integrates EukRep (West et al., 2018) for selecting the eukaryotic contigs from the initial assembly, the CONCOCT binning tool, GeneMark (Besemer et al., 2001) for prediction of eukaryotic genes, EukCC (Saary et al., 2020) for quality estimation from eukaryotic sequences, MAKER2 (Holt & Yandell, 2011) for gene annotation, and BUSCO (Waterhouse et al., 2017) for detection of single-copy orthologous genes. Lastly, in module 5, MuDoGeR groups the recovered MAGs into Operation Taxonomic Units (OTUs). Following, it maps the sequencing reads to the OTUs using two possible approaches: reduced or complete. The reduced method maps the recovered MAGs/UViGs/MABs on their respective libraries, while the complete method maps the recovered MAGs/UViGs/MABs on all available libraries. We kept the reduced mapping as default as, mostly, genomes recovered from different libraries may show coverage in a library even if, in reality, it was not present there. This is a consequence of the fact that a significant fraction of the genome encodes central metabolism genes that are highly conserved across most microbial species (Noor et al., 2010). Such genes are likely to map to a related species that may not be present in the sample. Later, it calculates the relative abundance and coverage tables from the mapped MAGs/UViGs/MABs and annotated prokaryotic genes within the assembled sequences. The user should have standardized results from a complete MAG pipeline by the end.

Finally, we designed MuDoGeR as a wrapper for several complex tools. It also makes all integrated software available for the user independently. This means that a more experienced user could integrate only pieces of MuDoGeR to their pipeline or even access a specific tool environment configured by MuDoGeR and use only the selected tool. Users can find a tutorial on activating these modules independently on the MuDoGeR GitHub page (https://github.com/mdsufz/MuDoGeR). Consequently, MuDoGeR modularity can give the researcher flexibility in their analysis and facilitate the investigator’s software management necessities.

MuDoGeR is designed to support Linux x64 systems. Some of the used software requires a large amount of RAM (e.g., GDTB-Tk, metaSPAdes). However, specific resource requirements vary depending on your data and its sequencing depth. For this reason, and to reduce the over/underuse of RAM, MuDoGeR attempts to calculate the amount of memory necessary for metaSPAdes.

The complete software installation requires approximately 170 GB, but MAKER2, from Module 4, uses 99 GB. The entire database requirements, considering all tools, are currently around 439.9 GB. In addition, it is recommended that the user provides multiple processing units since the analysis of several metagenomes simultaneously may require significant time. Consequently, the MuDoGeR Conda installation procedure allows it to be installed on high-performance computers or in cloud services such as <u>Amazon Elastic Compute Cloud</u>, <u>Google cloud</u>, or, for researchers in Germany, the <u>German Network for Bioinformatics Infrastructure</u>, allowing users to work with larger metagenome datasets.

## Results and discussion

We tested the MuDoGeR pipeline using 598 metagenome libraries (Illumina short reads) from public repositories (Supplementary Table 1). We selected these libraries to encompass metagenomes (574) and individual-isolated genomes (24). We specifically chose libraries from axenic genomes to test whether our pipeline can recover viruses in individual isolates (Supplementary Table 2). We could not evaluate our pipeline using a mock-community because, to the best of our knowledge, no standardized community containing prokaryotes, eukaryotes, and viruses existed by the time of our analysis. Only 8 out of the 574 metagenomic libraries showed no recovery of prokaryotic MAGs, putative viral contigs, or eukaryotic MABs (Supplementary Table 1). The following paragraphs detail the data for recovering prokaryotic MAGs, high-quality and complete UViGs, and eukaryotic MABs. Nevertheless, MuDoGeR can also generate output files containing information on ORFs. The accession numbers of the generated prokaryotic MAGs, high-quality and complete UViGs, and high-quality eukaryotic MABs can be found in Supplementary Tables 3, 4, and 5.

### Recovery of prokaryotes MAGs

MuDoGeR recovered 5726 prokaryotic MAGs encompassing 3850’ species level’ OTUs (Average Nucleotide Identity, ANI >0.95) (Supplementary Table 3) from the 574 metagenomic libraries used in this study. These included 1969 high-quality genomes (>90% completeness and <5% contamination) (Supplementary Table 3) (Figure 2A). These OTUs belonged to 3644 bacterial and 206 archaeal species, and GTDB-tk classified them over 77 Phyla, 141 Classes, 530 families, and 1110 genera (Supplementary Table 3). The number of prokaryotic MAGs per library ranged from 0 to 111 (average=9.98, standard deviation=15.81) (Supplementary Table 1) (Figure 2D).

**Figure 2.**
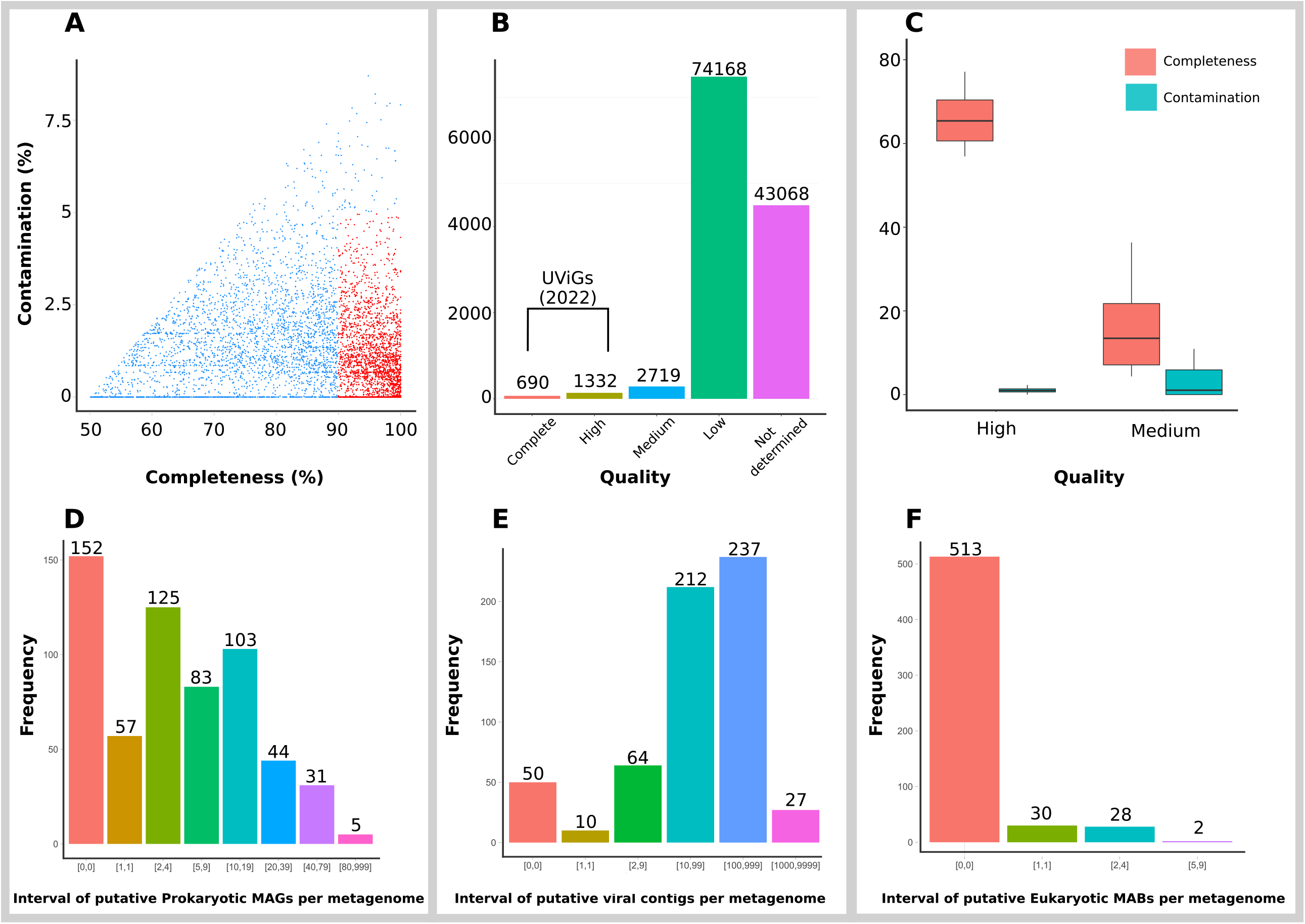
Results of prokaryotic and eukaryotic genome and viral contig recovery. **A –** Completeness and contamination of prokaryotic metagenome-assembled genomes (MAGs) with a quality score above 50 recovered from all libraries; **B –** Number of complete, high-quality, medium-quality, low-quality, and undetermined viral contigs (UViGs) recovered from all libraries; number on top of the bars indicate the number of UViGs per quality group.; **C –** Completeness and contamination boxplots for eukaryotic metagenome-assembled bins (MABs) recovered from all libraries; **D –** Frequency in the number of metagenomes recovering different numbers of MAGs; **E -** Frequency in the number of metagenomes recovering different numbers of putative viral contigs; **F -** Frequency in the number of metagenomes recovering different numbers of MABs. **D, E**, and **F** numbers between brackets indicate the intervals of MAGs, UViGs, and MABs, respectively; numbers on top of the bars indicate the number of metagenomes per interval.

MuDoGeR recovered no prokaryotic MAGs from approximately 27% (152) libraries. These libraries belonged to environments and materials with an extremely high diversity of microbes (e.g., soils) or samples where host DNA is present in higher yields than the microbial DNA before sequencing (e.g., endosymbiotic communities). Seven libraries showed more than 70 prokaryotic MAGs. From these, six metagenomes belonged to anaerobic reactors and one to a cyanobacterial mat (Supplementary Table 3). The relation between the sample’s diversity and complexity with the quality of recovered MAGs was previously observed in the work of (Bornemann et al., 2020). They identified that the bin quality of low complexity datasets (e.g., bioreactors) was significantly higher than in medium or high complexity datasets (e.g., soils).

### Recovery of UviGs

Our analysis uncovered 121 977 putative viral contigs (Supplementary Table 4). After dereplication and quality check, 2719 were classified as medium quality and 2022 as high-quality or complete UViGs (Supplementary Table 4) (Figure 2B). The number of dereplicated UViGs per library ranged from 0 to 166 (average=8.25 and standard deviation=15.40) (Supplementary Table 1) (Figure 2E). We observed at least one high-quality or complete UViG in approximately 85% (493) of the libraries. The libraries which did not yield any high-quality or complete UViGs belonged to environments and materials with high microbial diversity (e.g., soils) or samples where host DNA cannot be separated from microbial DNA before sequencing (e.g., endosymbiotic communities). Three libraries showed more than 100 high-quality or complete UVIGs. All of these belonged to aquatic environments (Supplementary Table 4). The high-quality and complete dereplicated UViGs belonged to the Order Caudovirales (978 UViGs) and Ligamenvirales (4 (UViGs). At the same time, most of them had their orders assigned as unclassified or undetermined (1092) by vConTACT2, indicating the high potential to identify new DNA viruses using MuDoGeR. Such a high number of unknown or unclassified viruses is not unusual when analyzing large (viral) metagenomic data sets (Santos-Medellin et al., 2021; Tisza & Buck, 2021).

### Recovery of Eukaryotic bins

We recovered a total of 52 eukaryotic MABs. Of these, 7 showed completeness above 40%, of which 5 showed less than 5% contamination (Figure 2C, Supplementary Table 5). As expected, due to the low number of MABs, we recovered no eukaryotes from 544 samples. The maximum number of MABs per sample was 6 (Figure 2F). The eukaryotic MABs showing more than 40% completeness were recovered from metagenomes from water and organic material retrieved from lakes, rivers, cropland, and urban samples (Supplementary Table 5). These bins are distributed over two phyla (Chlorophyta and Heterotrichea) and one species (*Micromonas commoda*) (Supplementary Table 5). Chlorophyta (i.e., green algae) are commonly found in marine habitats (Leliaert et al., 2012), which is consistent with the biome from which the samples were retrieved. Challenges in the recovery of eukaryotic MABs stem from the presence of repeat regions (Delmont & Eren, 2016), assembly of genomes using short reads (Pearman et al., 2020), a limited number of reference genomes (Pawlowski et al., 2012), and lack of software for predicting eukaryotic genes in entire metagenomes. Once the scientific community advances in eukaryotic genome reconstruction, we will add them to the MuDoGeR.

### Biodiversity analysis with MuDoGeR

Although we did not select metagenomes to perform biodiversity analysis with a specific research question in mind, we prepared the output of MuDoGeR to allow biodiversity analysis from prokaryotes, eukaryotes, and viruses alone or combined using standard biodiversity analyses with visualization pipelines (e.g., Phyloseq (McMurdie & Holmes, 2013)). To indicate its potential, we selected the samples with at least one eukaryotic bin and one medium, high-quality or complete UViG. Using these 18 libraries, we performed Beta diversity analyses using phyloseq to demonstrate the potential for genome-centric biodiversity analysis using MuDoGeR (Supplementary Figure 3).

## Conclusions

MuDoGeR is a user-friendly tool encompassing state-of-the-art software and pipelines able to recover prokaryotic, viral, and eukaryotic genomes from metagenomes in combination or individually. It extends to any study using Illumina short-sequence reads. Users can easily install all 20 tools and 50 custom scripts. MuDoGeR generates around 48 comprehensive files and folders containing the summary and parsing of the taxonomic information, quality estimation, genome annotation, coverage, and relative abundance calculation of the metagenome-assembled genomes. MuDoGeR requires only five simple commands to generate the output structure for the prokaryotic MAGs, UViGs, and eukaryotic MABs. Further, users may use any of the tools found in MuDoGeR by loading their specific Conda environment creating a flexible pipeline that expert users can adapt.

## Acknowledgments

This work was funded by the Deutsche Forschungsgemeinschaft (DFG, German Research Foundation) – project number 460129525. U.N.R, R.B.T., J.P.S, M.F.B., and F.B.C. were financed by the Helmholtz Young Investigator grant VH-NG-1248 Micro’ Big Data’. J.C.K was supported by the Fundação de Amparo à Pesquisa do Estado de São Paulo (FAPESP) [2019/03396-9] and [2022/03534-5]. J.P. was supported by Lundin Energy Norway AS within the framework of the GeneOil Project. This work also used the resources provided by the <u>German Network for Bioinformatics Infrastructure</u> (de.NBI)

<u>CLUE-TERRA consortium</u>: Alexander Bartholomäus, Alexandre Soares Rosado, Ana-Maria Fiore-Donno, André Carlos Ponce de Leon Ferreira de Carvalho, Cecile Gubry-Rangin, Daniel Machado, Danilo S. Sanches, Dirk Wagner, Gabriele Berg, Ines Mundic Mulec, Marie Muehe, Michael Bonkowski, Newton Gomes, Raquel Peixoto, Rodrigo Costa, Sabine Kleinsteuber, Simonetta Gribaldo, Tina Keller Costa, Vivian Pellizari.

## Data Availability Statement

MuDoGeR is open-source software available at https://github.com/mdsufz/MuDoGeR. All MAGs are available on NCBI SRA through https://www.ncbi.nlm.nih.gov/bioproject/PRJNA843551/. All high-quality and complete UViGs are available on the Helmholtz Center for Environmental Research - UFZ cloud services through the link https://nc.ufz.de/s/yFFBrceNoC6P3wY (password: pSNoAafzYB). All seven species-level MABs are available on NCBI SRA under the accessions SAMN26329102, SAMN26244053, SAMN26329113, SAMN26302841, SAMN26302842, SAMN26329290, and SAMN26302904.

## Conflict of interests

The authors declare that they have no competing interests.

## Authors’ contributions

UNR conceptualized, coordinated, and supervised all work. RK joined the conceptualization of the viral genome recovery. JPS and PB joined the conceptualization of the eukaryotic genome recovery. PS joined the conceptualization for reaching non-bioinformaticians and user-friendliness of the pipeline. JCK, RBT, MFB, FBC, LMVB, and MNB worked on writing the code for the pipeline and its GitHub page. EF helped with the GitHub and standardization of the metadata from the metagenomes and genomes used in the study. JP and AP tested the tool for its user-friendly aspect and improved the GitHub and code. PFS discussed the conceptualization and the user-friendly aspect of the tool. All authors helped in the writing of the manuscript. The CLUE-TERRA consortium was instrumental in discussing and designing the potential functionality of MuDoGeR and its user-friendly aspect.

## Notes

https://github.com/mdsufz/MuDoGeR

### Competing Interest Statement

The authors have declared no competing interest.

### Summary of Updates

We improved the flow in the text and added a comparison of MuDoGeR to other tools performing the recovery of genomes from metagenomes.

## References

Albertsen, M., Hugenholtz, P., Skarshewski, A., Nielsen, K. L., Tyson, G. W., & Nielsen, P. H. (2013). Genome sequences of rare, uncultured bacteria obtained by differential coverage binning of multiple metagenomes. Nature Biotechnology, 31(6), 533–538. https://doi.org/10.1038/nbt.2579

Alneberg, J., Bjarnason, B. S., Bruijn, I. de, Schirmer, M., Quick, J., Ijaz, U. Z., Lahti, L., Loman, N. J., Andersson, A. F., & Quince, C. (2014). Binning metagenomic contigs by coverage and composition. Nature Methods, 11(11), 1144–1146. https://doi.org/10.1038/nmeth.3103

Besemer, J., Lomsadze, A., & Borodovsky, M. (2001). GeneMarkS: A self-training method for prediction of gene starts in microbial genomes. Implications for finding sequence motifs in regulatory regions. Nucleic Acids Research, 29(12), 2607–2618.

Bin Jang, H., Bolduc, B., Zablocki, O., Kuhn, J. H., Roux, S., Adriaenssens, E. M., Brister, J. R., Kropinski, A. M., Krupovic, M., Lavigne, R., Turner, D., & Sullivan, M. B. (2019). Taxonomic assignment of uncultivated prokaryotic virus genomes is enabled by gene-sharing networks. Nature Biotechnology, 37(6), 632–639. https://doi.org/10.1038/s41587-019-0100-8

Bornemann, T. L. V., Esser, S. P., Stach, T. L., Burg, T., & Probst, A. J. (2020). UBin – a manual refining tool for metagenomic bins designed for educational purposes. BioRxiv, 2020.07.15.204776. https://doi.org/10.1101/2020.07.15.204776

Chaumeil, P.-A., Mussig, A. J., Hugenholtz, P., & Parks, D. H. (2020). GTDB-Tk: A toolkit to classify genomes with the Genome Taxonomy Database. Bioinformatics, 36(6), 1925–1927. https://doi.org/10.1093/bioinformatics/btz848

Corrêa, F. B., Saraiva, J. P., Stadler, P. F., & da Rocha, U. N. (2020). TerrestrialMetagenomeDB: A public repository of curated and standardized metadata for terrestrial metagenomes. Nucleic Acids Research, 48(D1), D626–D632. https://doi.org/10.1093/nar/gkz994

Delmont, T. O., & Eren, A. M. (2016). Identifying contamination with advanced visualization and analysis practices: Metagenomic approaches for eukaryotic genome assemblies. PeerJ, 4, e1839. https://doi.org/10.7717/peerj.1839

Dias, O., Saraiva, J., Faria, C., Ramirez, M., Pinto, F., & Rocha, I. (2019). IDS372, a Phenotypically Reconciled Model for the Metabolism of Streptococcus pneumoniae Strain R6. Frontiers in Microbiology, 10, 1283. https://doi.org/10.3389/fmicb.2019.01283

Evans Paul N., Parks Donovan H., Chadwick Grayson L., Robbins Steven J., Orphan Victoria J., Golding Suzanne D., & Tyson Gene W. (2015). Methane metabolism in the archaeal phylum Bathyarchaeota revealed by genome-centric metagenomics. Science, 350(6259), 434–438. https://doi.org/10.1126/science.aac7745

Galiez, C., Siebert, M., Enault, F., Vincent, J., & Söding, J. (2017). WIsH: Who is the host? Predicting prokaryotic hosts from metagenomic phage contigs. Bioinformatics, 33(19), 3113–3114. https://doi.org/10.1093/bioinformatics/btx383

Grüning, B., Dale, R., Sjödin, A., Chapman, B. A., Rowe, J., Tomkins-Tinch, C. H., Valieris, R., & Köster, J. (2018). Bioconda: Sustainable and comprehensive software distribution for the life sciences. Nature Methods, 15(7), 475–476. https://doi.org/10.1038/s41592-018-0046-7

Guo, J., Bolduc, B., Zayed, A. A., Varsani, A., Dominguez-Huerta, G., Delmont, T. O., Pratama, A. A., Gazitúa, M. C., Vik, D., Sullivan, M. B., & Roux, S. (2021). VirSorter2: A multi-classifier, expert-guided approach to detect diverse DNA and RNA viruses. Microbiome, 9(1), 37. https://doi.org/10.1186/s40168-020-00990-y

Holt, C., & Yandell, M. (2011). MAKER2: An annotation pipeline and genome-database management tool for second-generation genome projects. BMC Bioinformatics, 12(1), 491. https://doi.org/10.1186/1471-2105-12-491

Kallies, R., Hölzer, M., Brizola Toscan, R., Nunes da Rocha, U., Anders, J., Marz, M., & Chatzinotas, A. (2019). Evaluation of Sequencing Library Preparation Protocols for Viral Metagenomic Analysis from Pristine Aquifer Groundwaters. Viruses, 11(6), 484. https://doi.org/10.3390/v11060484

Kang, D. D., Froula, J., Egan, R., & Wang, Z. (2015). MetaBAT, an efficient tool for accurately reconstructing single genomes from complex microbial communities. PeerJ, 3, e1165. https://doi.org/10.7717/peerj.1165

Kasmanas, J. C., Bartholomäus, A., Corrêa, F. B., Tal, T., Jehmlich, N., Herberth, G., von Bergen, M., Stadler, P. F., Carvalho, A. C. P. de L. F. de, & Nunes da Rocha, U. (2021). HumanMetagenomeDB: A public repository of curated and standardized metadata for human metagenomes. Nucleic Acids Research, 49(D1), D743–D750. https://doi.org/10.1093/nar/gkaa1031

Keller-Costa, T., Lago-Lestón, A., Saraiva, J. P., Toscan, R., Silva, S. G., Gonçalves, J., Cox, C. J., Kyrpides, N., Nunes da Rocha, U., & Costa, R. (2021). Metagenomic insights into the taxonomy, function, and dysbiosis of prokaryotic communities in octocorals. Microbiome, 9(1), 72. https://doi.org/10.1186/s40168-021-01031-y

Kieft, K., Zhou, Z., & Anantharaman, K. (2020). VIBRANT: Automated recovery, annotation and curation of microbial viruses, and evaluation of viral community function from genomic sequences. Microbiome, 8(1), 90. https://doi.org/10.1186/s40168-020-00867-0

Koren, S., Treangen, T. J., & Pop, M. (2011). Bambus 2: Scaffolding metagenomes. Bioinformatics, 27(21), 2964–2971. https://doi.org/10.1093/bioinformatics/btr520

Leliaert, F., Smith, D. R., Moreau, H., Herron, M. D., Verbruggen, H., Delwiche, C. F., & De Clerck, O. (2012). Phylogeny and Molecular Evolution of the Green Algae. Critical Reviews in Plant Sciences, 31(1), 1–46. https://doi.org/10.1080/07352689.2011.615705

Li, D., Luo, R., Liu, C.-M., Leung, C.-M., Ting, H.-F., Sadakane, K., Yamashita, H., & Lam, T.-W. (2016). MEGAHIT v1.0: A fast and scalable metagenome assembler driven by advanced methodologies and community practices. Methods (San Diego, Calif.), 102, 3–11. https://doi.org/10.1016/j.ymeth.2016.02.020

Liu, B., Sträuber, H., Saraiva, J., Harms, H., Silva, S. G., Kasmanas, J. C., Kleinsteuber, S., & Nunes da Rocha, U. (2022). Machine learning-assisted identification of bioindicators predicts medium-chain carboxylate production performance of an anaerobic mixed culture. Microbiome, 10(1), 48. https://doi.org/10.1186/s40168-021-01219-2

López-Mondéjar, R., Tláskal, V., Větrovský, T., Štursová, M., Toscan, R., Nunes da Rocha, U., & Baldrian, P. (2020). Metagenomics and stable isotope probing reveal the complementary contribution of fungal and bacterial communities in the recycling of dead biomass in forest soil. Soil Biology and Biochemistry, 148, 107875. https://doi.org/10.1016/j.soilbio.2020.107875

Mangul, S., Mosqueiro, T., Abdill, R. J., Duong, D., Mitchell, K., Sarwal, V., Hill, B., Brito, J., Littman, R. J., Statz, B., Lam, A. K.-M., Dayama, G., Grieneisen, L., Martin, L. S., Flint, J., Eskin, E., & Blekhman, R. (2019). Challenges and recommendations to improve the installability and archival stability of omics computational tools. PLOS Biology, 17(6), e3000333. https://doi.org/10.1371/journal.pbio.3000333

McMurdie, P. J., & Holmes, S. (2013). phyloseq: An R Package for Reproducible Interactive Analysis and Graphics of Microbiome Census Data. PLOS ONE, 8(4), e61217. https://doi.org/10.1371/journal.pone.0061217

Melkonian, C., Fillinger, L., Atashgahi, S., da Rocha, U. N., Kuiper, E., Olivier, B., Braster, M., Gottstein, W., Helmus, R., Parsons, J. R., Smidt, H., van der Waals, M., Gerritse, J., Brandt, B. W., Röling, W. F. M., Molenaar, D., & van Spanning, R. J. M. (2021). High biodiversity in a benzene-degrading nitrate-reducing culture is sustained by a few primary consumers. Communications Biology, 4(1), 1–12. https://doi.org/10.1038/s42003-021-01948-y

Namiki, T., Hachiya, T., Tanaka, H., & Sakakibara, Y. (2012). MetaVelvet: An extension of Velvet assembler to de novo metagenome assembly from short sequence reads. Nucleic Acids Research, 40(20), e155–e155. https://doi.org/10.1093/nar/gks678

Nayfach, S., Camargo, A. P., Schulz, F., Eloe-Fadrosh, E., Roux, S., & Kyrpides, N. C. (2021). CheckV assesses the quality and completeness of metagenome-assembled viral genomes. Nature Biotechnology, 39(5), 578–585. https://doi.org/10.1038/s41587-020-00774-7

Noor, E., Eden, E., Milo, R., & Alon, U. (2010). Central Carbon Metabolism as a Minimal Biochemical Walk between Precursors for Biomass and Energy. Molecular Cell, 39(5), 809–820. https://doi.org/10.1016/j.molcel.2010.08.031

Nurk, S., Meleshko, D., Korobeynikov, A., & Pevzner, P. A. (2017). metaSPAdes: A new versatile metagenomic assembler. Genome Research, 27(5), 824–834. https://doi.org/10.1101/gr.213959.116

Oliveira Monteiro, L. M., Saraiva, J. P., Brizola Toscan, R., Stadler, P. F., Silva-Rocha, R., & Nunes da Rocha, U. (2022). PredicTF: Prediction of bacterial transcription factors in complex microbial communities using deep learning. Environmental Microbiome, 17(1), 7. https://doi.org/10.1186/s40793-021-00394-x

Parks, D. H., Imelfort, M., Skennerton, C. T., Hugenholtz, P., & Tyson, G. W. (2015). CheckM: Assessing the quality of microbial genomes recovered from isolates, single cells, and metagenomes. Genome Research, 25(7), 1043–1055. https://doi.org/10.1101/gr.186072.114

Parks, D. H., Rinke, C., Chuvochina, M., Chaumeil, P.-A., Woodcroft, B. J., Evans, P. N., Hugenholtz, P., & Tyson, G. W. (2017). Recovery of nearly 8,000 metagenome-assembled genomes substantially expands the tree of life. Nature Microbiology, 2(11), 1533–1542. https://doi.org/10.1038/s41564-017-0012-7

Pawlowski, J., Audic, S., Adl, S., Bass, D., Belbahri, L., Berney, C., Bowser, S. S., Cepicka, I., Decelle, J., Dunthorn, M., Fiore-Donno, A. M., Gile, G. H., Holzmann, M., Jahn, R., Jirků, M., Keeling, P. J., Kostka, M., Kudryavtsev, A., Lara, E., … Vargas, C. de. (2012). CBOL Protist Working Group: Barcoding Eukaryotic Richness beyond the Animal, Plant, and Fungal Kingdoms. PLOS Biology, 10(11), e1001419. https://doi.org/10.1371/journal.pbio.1001419

Pearman, W. S., Freed, N. E., & Silander, O. K. (2020). Testing the advantages and disadvantages of short- and long-read eukaryotic metagenomics using simulated reads. BMC Bioinformatics, 21(1), 220. https://doi.org/10.1186/s12859-020-3528-4

Peng, Y., Leung, H. C. M., Yiu, S. M., & Chin, F. Y. L. (2012). IDBA-UD: A de novo assembler for single-cell and metagenomic sequencing data with highly uneven depth. Bioinformatics, 28(11), 1420–1428. https://doi.org/10.1093/bioinformatics/bts174

Ren, J., Ahlgren, N. A., Lu, Y. Y., Fuhrman, J. A., & Sun, F. (2017). VirFinder: A novel k-mer based tool for identifying viral sequences from assembled metagenomic data. Microbiome, 5(1), 69. https://doi.org/10.1186/s40168-017-0283-5

Saary, P., Mitchell, A. L., & Finn, R. D. (2020). Estimating the quality of eukaryotic genomes recovered from metagenomic analysis with EukCC. Genome Biology, 21(1), 244. https://doi.org/10.1186/s13059-020-02155-4

Santos-Medellin, C., Zinke, L. A., ter Horst, A. M., Gelardi, D. L., Parikh, S. J., & Emerson, J. B. (2021). Viromes outperform total metagenomes in revealing the spatiotemporal patterns of agricultural soil viral communities. The ISME Journal, 15(7), 1956–1970. https://doi.org/10.1038/s41396-021-00897-y

Saraiva, J. P., Bartholomäus, A., Kallies, R., Gomes, M., Bicalho, M., Coelho Kasmanas, J., Vogt, C., Chatzinotas, A., Stadler, P., Dias, O., & Nunes da Rocha, U. (2021). OrtSuite: From genomes to prediction of microbial interactions within targeted ecosystem processes. Life Science Alliance, 4(12), e202101167. https://doi.org/10.26508/lsa.202101167

Saraiva, J. P. L. F., Zubiria-Barrera, C., Klassert, T. E., Lautenbach, M. J., Blaess, M., Claus, R. A., Slevogt, H., & König, R. (2017). Combination of classifiers identifies fungal-specific activation of lysosome genes in human monocytes. Frontiers in Microbiology, 8(NOV). Scopus. https://doi.org/10.3389/fmicb.2017.02366

Saraiva, J. P., Oswald, M., Biering, A., Röll, D., Assmann, C., Klassert, T., Blaess, M., Czakai, K., Claus, R., Löffler, J., & others. (2017). Fungal biomarker discovery by integration of classifiers. BMC Genomics, 18(1), 601.

Saraiva, J. P., Worrich, A., Karakoç, C., Kallies, R., Chatzinotas, A., Centler, F., & Nunes da Rocha, U. (2021). Mining Synergistic Microbial Interactions: A Roadmap on How to Integrate Multi-Omics Data. Microorganisms, 9(4), 840. https://doi.org/10.3390/microorganisms9040840

Seemann, T. (2014). Prokka: Rapid prokaryotic genome annotation. Bioinformatics (Oxford, England), 30(14), 2068–2069. https://doi.org/10.1093/bioinformatics/btu153

Sharon, I., Morowitz, M. J., Thomas, B. C., Costello, E. K., Relman, D. A., & Banfield, J. F. (2013). Time series community genomics analysis reveals rapid shifts in bacterial species, strains, and phage during infant gut colonization. Genome Research, 23(1), 111–120. https://doi.org/10.1101/gr.142315.112

Sieber, C. M. K., Probst, A. J., Sharrar, A., Thomas, B. C., Hess, M., Tringe, S. G., & Banfield, J. F. (2018). Recovery of genomes from metagenomes via a dereplication, aggregation and scoring strategy. Nature Microbiology, 3(7), 836–843. https://doi.org/10.1038/s41564-018-0171-1

Tisza, M. J., & Buck, C. B. (2021). A catalog of tens of thousands of viruses from human metagenomes reveals hidden associations with chronic diseases. Proceedings of the National Academy of Sciences, 118(23), e2023202118. https://doi.org/10.1073/pnas.2023202118

Tláskal, V., Brabcová, V., Větrovský, T., Jomura, M., López-Mondéjar, R., Monteiro, L. M. O., Saraiva, J. P., Human, Z. R., Cajthaml, T., Rocha, U. N. da, & Baldrian, P. (2021). Complementary Roles of Wood-Inhabiting Fungi and Bacteria Facilitate Deadwood Decomposition. MSystems, 6(1). https://doi.org/10.1128/mSystems.01078-20

Tyson, G. W., Chapman, J., Hugenholtz, P., Allen, E. E., Ram, R. J., Richardson, P. M., Solovyev, V. V., Rubin, E. M., Rokhsar, D. S., & Banfield, J. F. (2004). Community structure and metabolism through reconstruction of microbial genomes from the environment. Nature, 428(6978), 37–43. https://doi.org/10.1038/nature02340

Uritskiy, G. V., DiRuggiero, J., & Taylor, J. (2018). MetaWRAP—a flexible pipeline for genome-resolved metagenomic data analysis. Microbiome, 6(1), 158. https://doi.org/10.1186/s40168-018-0541-1

Waterhouse, R. M., Seppey, M., Simão, F. A., Manni, M., Ioannidis, P., Klioutchnikov, G., Kriventseva, E. V., & Zdobnov, E. M. (2017). BUSCO applications from quality assessments to gene prediction and phylogenomics. Molecular Biology and Evolution. https://doi.org/10.1093/molbev/msx319

West, P. T., Probst, A. J., Grigoriev, I. V., Thomas, B. C., & Banfield, J. F. (2018). Genome-reconstruction for eukaryotes from complex natural microbial communities. Genome Research, 28(4), 569–580. https://doi.org/10.1101/gr.228429.117

Wilkinson, M. D., Dumontier, M., Aalbersberg, Ij. J., Appleton, G., Axton, M., Baak, A., Blomberg, N., Boiten, J.-W., da Silva Santos, L. B., Bourne, P. E., Bouwman, J., Brookes, A. J., Clark, T., Crosas, M., Dillo, I., Dumon, O., Edmunds, S., Evelo, C. T., Finkers, R., … Mons, B. (2016). The FAIR Guiding Principles for scientific data management and stewardship. Scientific Data, 3(1), 160018. https://doi.org/10.1038/sdata.2016.18

Wooley, J. C., & Ye, Y. (2010). Metagenomics: Facts and Artifacts, and Computational Challenges. Journal of Computer Science and Technology, 25(1), 71–81. https://doi.org/10.1007/s11390-010-9306-4

Wu, Y.-W., Simmons, B. A., & Singer, S. W. (2016). MaxBin 2.0: An automated binning algorithm to recover genomes from multiple metagenomic datasets. Bioinformatics (Oxford, England), 32(4), 605–607. https://doi.org/10.1093/bioinformatics/btv638

