## Supplementary Figure 2 for "MuDoGeR: Multi-Domain Genome Recovery from metagenomes made easy"

[1]Department of Environmental Microbiology, Helmholtz Centre for Environmental Research – UFZ, Leipzig, Germany.
[2]Institute of Mathematics and Computer Sciences, University of São Paulo, São Carlos, Brazil.
[3] Biotechnical Faculty, University of Ljubljana, Ljubljana, Slovenia.
[4]Environmental Microbiology and Biotechnology, Department of Chemistry, University of Duisburg-Essen, 45141, Essen, Germany.
[5]Laboratory of Environmental Microbiology, Institute of Microbiology of the Czech Academy of Sciences, Videnska 1083, Praha 4, 14220, Czech Republic.
[6]Department of Computer Science and Interdisciplinary Center of Bioinformatics, University of Leipzig, Leipzig, Germany. Max Planck Institute for Mathematics in the Sciences, Leipzig, Germany. Institute for Theoretical Chemistry, Univ. of Vienna, Austria. The Santa Fe Institute, Santa Fe, NM.

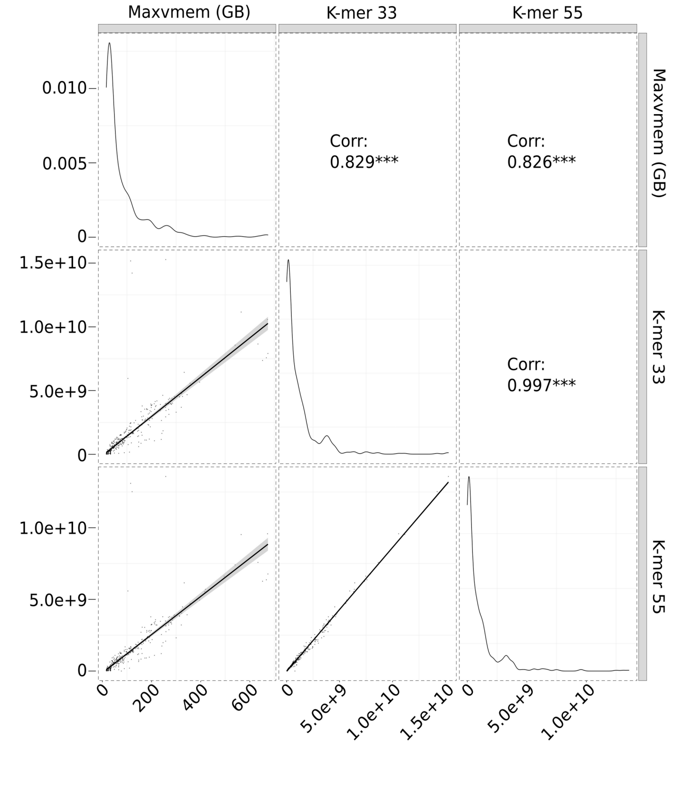


**Supplementary Figure 2 –** Correlation analysis of total Random Access Memory (RAM) and chimer frequencies.
