## Supplementary Figure 3 for "MuDoGeR: Multi-Domain Genome Recovery from metagenomes made easy"

[1]Department of Environmental Microbiology, Helmholtz Centre for Environmental Research – UFZ, Leipzig, Germany.
[2]Institute of Mathematics and Computer Sciences, University of São Paulo, São Carlos, Brazil.
[3] Biotechnical Faculty, University of Ljubljana, Ljubljana, Slovenia.
[4]Environmental Microbiology and Biotechnology, Department of Chemistry, University of Duisburg-Essen, 45141, Essen, Germany.
[5]Laboratory of Environmental Microbiology, Institute of Microbiology of the Czech Academy of Sciences, Videnska 1083, Praha 4, 14220, Czech Republic.
[6]Department of Computer Science and Interdisciplinary Center of Bioinformatics, University of Leipzig, Leipzig, Germany. Max Planck Institute for Mathematics in the Sciences, Leipzig, Germany. Institute for Theoretical Chemistry, Univ. of Vienna, Austria. The Santa Fe Institute, Santa Fe, NM.

**A**

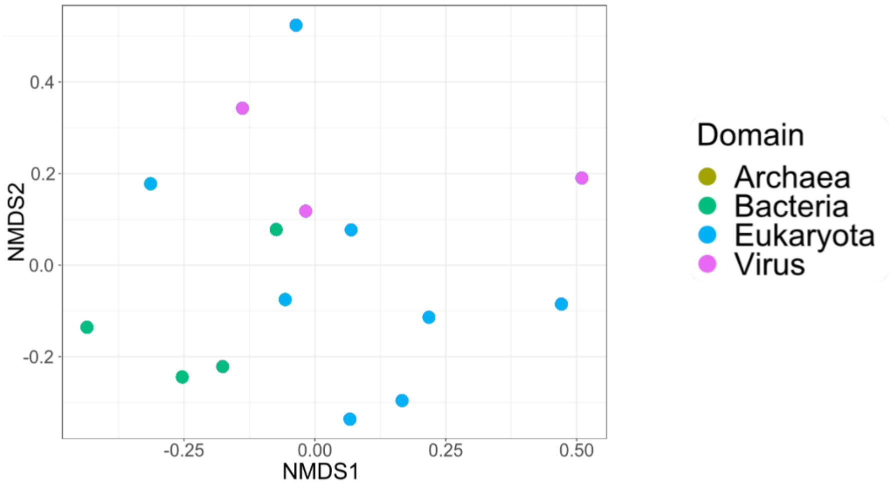


**B**

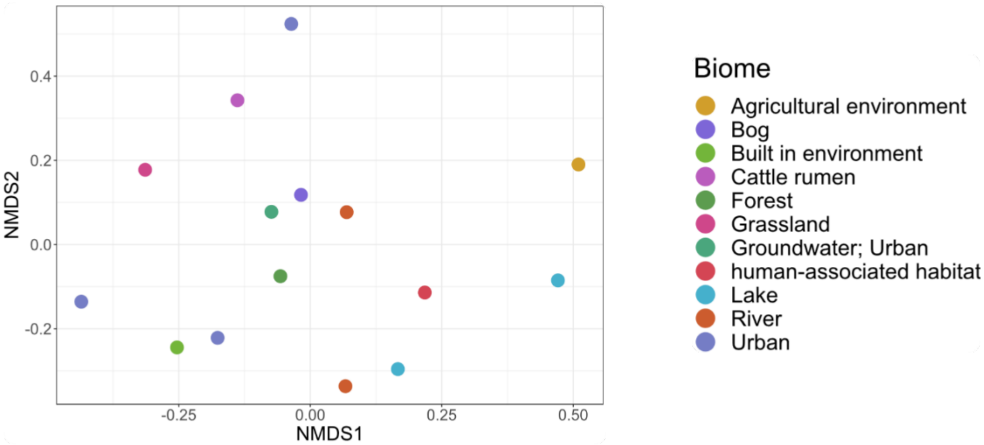


**Supplementary Figure 3 – Beta diversity analysis.** A – NMDS plot containing only the samples where prokaryotes, viruses, and eukaryotes genomes were recovered. The figure was created using the plot_ordination function from phyloseq using the relative abundance from the recovered genomes as calculated by MuDoGeR and colored by Biome;

B – NMDS plot containing only the samples where prokaryotes, viruses, and eukaryotes genomes were recovered. The figure was created using the plot_ordination function from phyloseq with type set to 'taxa'. The plot was created using the relative abundance from the recovered genomes grouped by their lowest level and colored by the taxonomic domain. This figure shows a potential separation from the samples according to their Domain composition recovered.
